# Low-heteroplasmy mitochondrial DNA mutations improve clonal reconstruction of human cells

**DOI:** 10.64898/2026.08.17.745291

**Authors:** Chen Weng, Teng Gao, William Colgan, Isaac Johnson, Jonas Gudera, Michael Poeschla, Jonathan S. Weissman, Vijay G. Sankaran

## Abstract

Reconstructing clonal relationships among human cells is fundamental to understanding development, aging, and disease. Somatic mitochondrial DNA (mtDNA) mutations act as endogenous single-cell barcodes measurable alongside cell-state profiles, but lineage tracing has traditionally focused on high-heteroplasmy variants, which are easier to detect but few and potentially shaped by selection. Whether the more abundant lower-heteroplasmy variants encode bona fide lineage information has not been tested against an independent clonal reference. Using lentiviral barcoding of human hematopoietic cells to establish ground-truth clone identities, we show that after stringent molecule-level error filtering, mutation calls below 10% per-cell heteroplasmy account for roughly half of all lineage-informative calls. Retaining the full heteroplasmy spectrum approximately doubled the clonal-assignment area under the precision– recall curve relative to a >10% cutoff, and single-molecule-supported calls improved recovery when retained collectively. These findings establish lower-heteroplasmy mtDNA mutations as an abundant, bona fide record of clonal history, substantially expanding the clonal resolution attainable in human tissues without genetic engineering.

## Background

Endogenous lineage recorders are especially valuable in human tissues because they capture clonal history without prior genetic engineering. Among the available classes, somatic mtDNA mutations are rich in lineage information, scalable to hundreds of thousands of cells, and measurable jointly with transcriptomic and chromatin-accessibility readouts of the same cell [1– 4]. The mitochondrial genome is present in hundreds to thousands of copies per cell, therefore a mutation typically occupies only a fraction. This fraction, the heteroplasmy level, ranges from 0 to 100% [5]. mtDNA has a ~10–20-fold higher mutation rate than nuclear DNA, generating a large and diverse pool of somatic mutations, with most occurring at low heteroplasmy [6–8]. However, owing to the challenges of mtDNA mutation detection, single-cell mtDNA lineage-tracing studies have traditionally focused on high-heteroplasmy somatic variants (e.g. >10%), which are easier to detect but few and thus capture only a limited proportion of the available lineage information. The more numerous lower-heteroplasmy variants can substantially expand the pool of somatic lineage markers [3,5,6,9], but their detection in single cells is more challenging, as each observation is supported by fewer mutant molecules per cell. We previously developed ReDeeM, which implemented single-molecule consensus correction with endogenous unique molecular identifiers (eUMIs), mitigating PCR/sequencing derived artifacts and enabling high-confidence detection of >10-fold more somatic mtDNA mutations [3,10]. Yet whether these lower-heteroplasmy observations encode bona fide lineage information has not been tested against an independent ground-truth clonal reference [11]. Here we address this using lentiviral barcoding of human hematopoietic stem and progenitor cells, which assigns each founder cell an independent ground-truth clonal identity. We show that mutation calls below 10% heteroplasmy carry more than half of the usable lineage information and substantially improve recovery of true clonal relationships, expanding what naturally occurring mitochondrial variation can reveal about human cell history.

## Results and discussion

We first asked whether variant calls with low molecule counts per cell retain molecular hallmarks of genuine mtDNA mutations. Beyond eUMI consensus correction, we added post-consensus filtering in redeemR2.0 (**Fig. 1A**): 9-bp fragment-end trimming to remove residual edge-enriched artifacts; a per-variant binomial goodness-of-fit test retaining variants whose cell-to-cell distributions deviate from random technical noise; and removal of germline-like variants by population frequency (MITOMAP; ref. 12). After filtering, calls supported by one, two, or ≥3 eUMIs per cell all showed the transition-dominated signature of bona fide mtDNA mutations, with low transversion ratios (~0.03; true-signal range 0.03–0.1 versus ~0.65 background) and uniform distribution along read fragments (**Fig. 1B**). Importantly, these refinements cost few variant numbers: in a human hematopoietic stem cell dataset, 3,932 of 4,394 called variants (89.5%) passed the additional filters. The resulting set still contains >10-fold more confident variants than prior methods lacking consensus-corrections (10) (**Fig. 1C**).

**Fig. 1.**
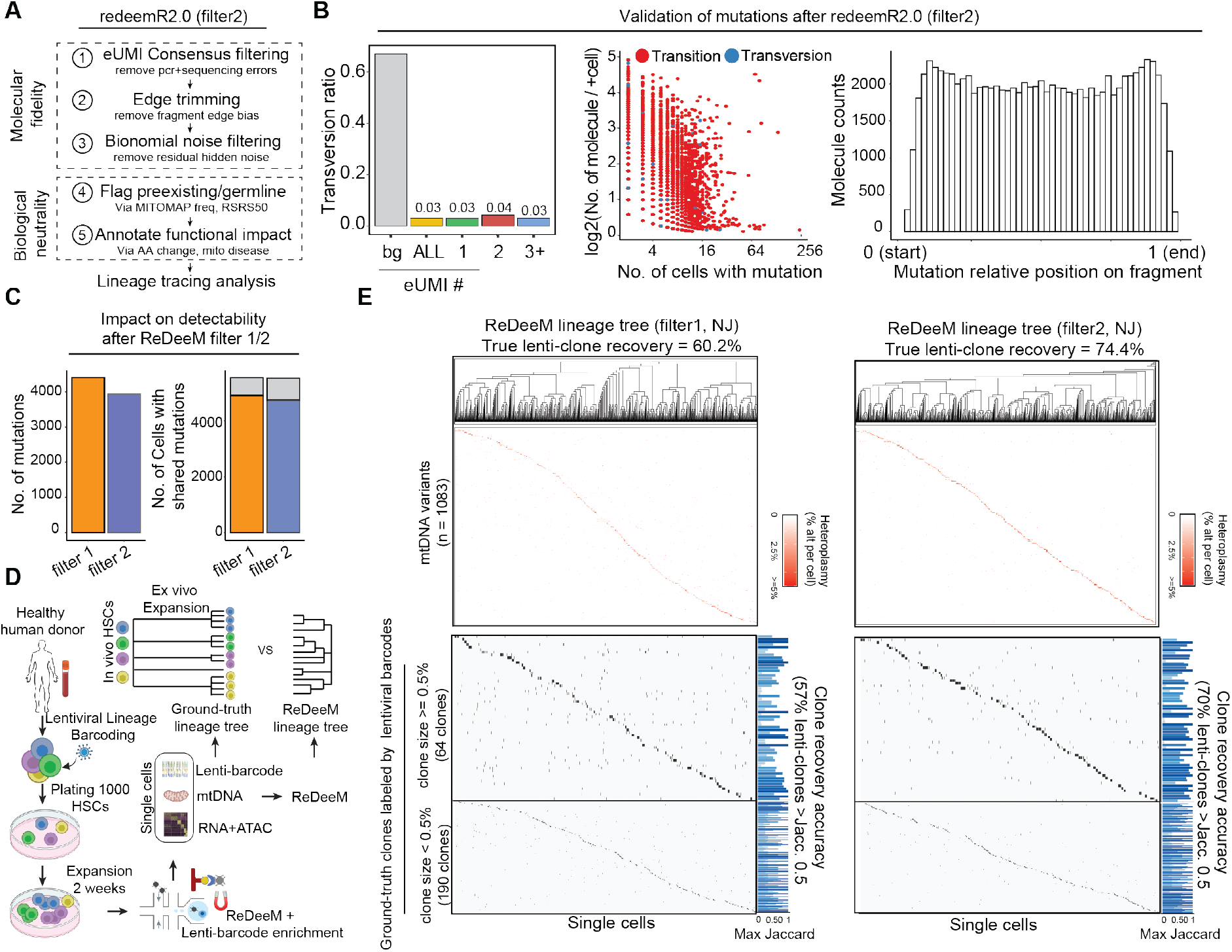
redeemR2.0 filtering and ground-truth–anchored benchmarking of mtDNA lineage tracing. **(A)** filter2 workflow: eUMI consensus filtering, fragment-edge trimming, binomial noise filtering, and variant annotation. **(B)** Variants retained after filter2 (Young1-HSC; ref. 3): transversion ratio by molecular support (Left), mutation frequency (Center), and fragment position (Right). **(C)** Variants removed by filter2 relative to filter1. **(D)** Ground-truth design: LARRY-barcoded primary human HSCs expanded ex vivo and profiled by ReDeeM with paired RNA, ATAC, mtDNA, and barcode recovery. **(E)** ReDeeM trees from filter1 or filter2 versus lenti-clones. Cells ordered by tree; heteroplasmy heatmaps above lenti-clone assignments; bars, maximum Jaccard similarity per lenti-clone.

To benchmark mtDNA-based lineage inference, we used a dataset in which primary human HSCs were labeled with lentiviral lineage and RNA-recovery barcodes (LARRY v2), expanded and differentiated ex vivo, then profiled by ReDeeM with simultaneous mtDNA mutation and lentiviral barcode recovery [11]. Cells sharing a lentiviral barcode define a lenti-clone, and the 254 recovered lenti-clones served as ground truth (**Fig. 1D**). ReDeeM reconstructs lineage from cell-cell mtDNA variant profile similarity. We scored a lenti-clone as recovered when its best-matching mtDNA-inferred clade shared >0.5 Jaccard similarity in cell membership. Trees built with the original strategy (filter1; ref. 3) or with redeemR2.0 (filter2) both recovered lenti-clones effectively, with filter2 improving recovery from 60.2% to 74.4% (**Fig. 1E**).

We next assessed how variant selection, meaning filtering strategy and heteroplasmy cutoffs, affects clonal tracing (**Fig. 2A**). We used MitoDrift as our lineage inference framework [11], which provides confidence-aware clade reconstruction from mtDNA variants, enabling us to assess both true-clone recovery and clade precision. We defined precision as the fraction of inferred clades that matched a ground-truth clone above a given Jaccard threshold, and recall as the fraction of ground-truth clones recovered by an inferred clade above that threshold. Benchmarking identified redeemR2.0 (filter2) as the best strategy for downstream lineage reconstruction: its precision– recall curve dominated all others (**Figure 2B**). Both filter1 and filter2 outperformed mgatk-like settings, which lack UMI-based error correction, as well as more stringent heteroplasmy cutoffs. Retaining the full filter2 set raised the area under the precision–recall curve (AUPRC) from 0.322 to 0.640 at Jaccard ≥0.5 relative to a >10% heteroplasmy restriction, and from 0.203 to 0.433 at Jaccard ≥0.75, gains of 2- and 2.1-fold. Including 1-eUMI variants also helped, raising AUPRC from 0.573 to 0.640 and from 0.356 to 0.433 at the two thresholds, indicating this class contributes more lineage signal than noise. We also observed a precision ceiling: under aggressive heteroplasmy filtering, precision did not improve proportionally while recall declined.

**Fig. 2.**
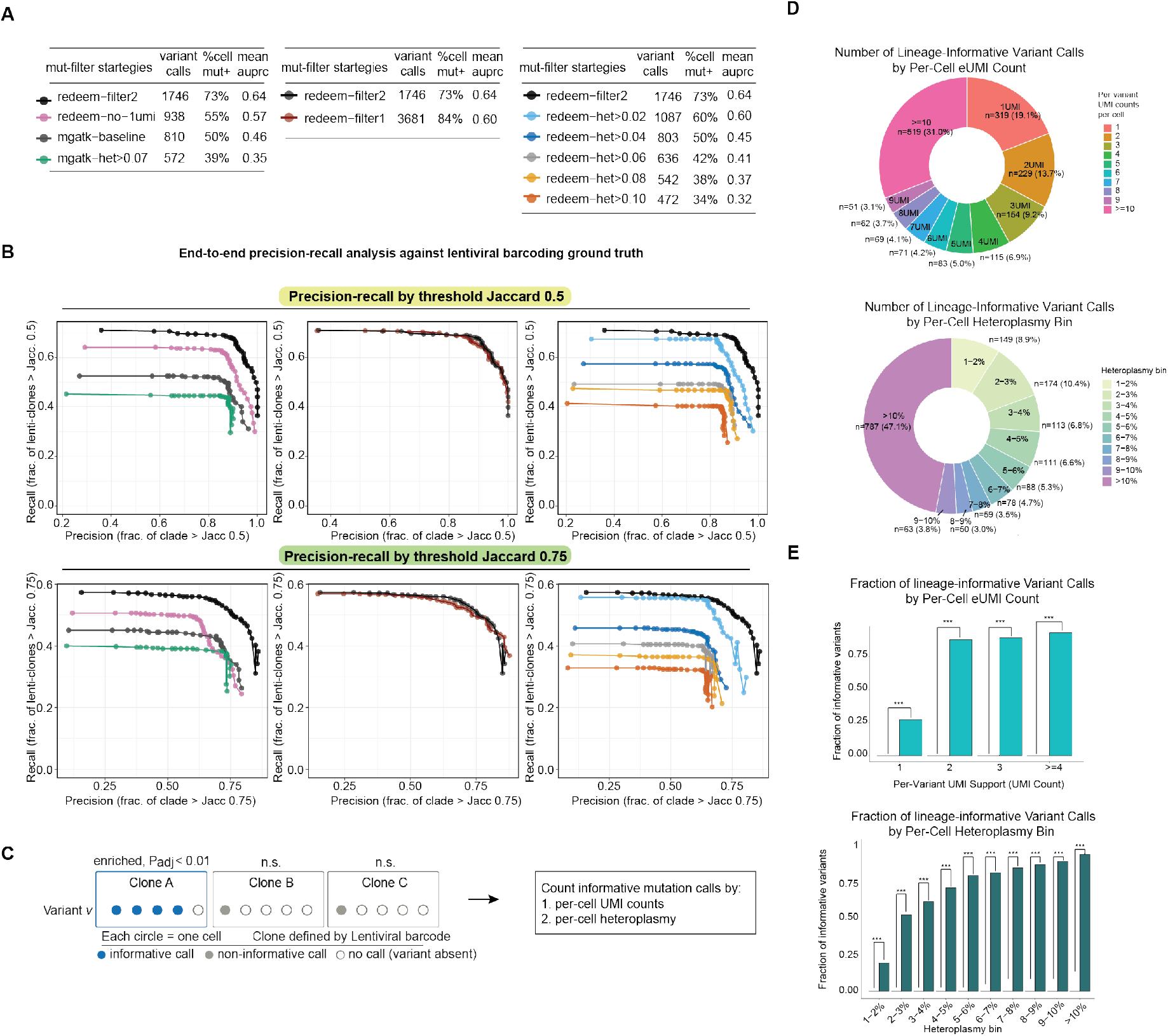
Benchmarking mtDNA lineage tracing across filtering strategies and heteroplasmy thresholds. **(A)** Strategies evaluated, with variant-call numbers, percentage of cells with ≥1 mutation, and mean AUPRC. **(B)** End-to-end precision–recall against lenti-clones across MitoDrift clade-support thresholds, at Jaccard cutoffs of 0.5 and 0.75. **(C)** Definition of lineage-informative calls by variant enrichment within lenti-clones. **(D)** Contribution of lineage-informative calls by eUMI support and per-cell heteroplasmy. **(E)** Fraction of lineage-informative calls within each group versus randomized background.

We next asked how lineage information is distributed across individual variants. Using lenti-clones as ground truth, we tested each variant for enrichment within individual clones (Fisher’s exact test, FDR-corrected across clones). We term a mutation call (the detection of a given variant in a given cell) lineage-informative when the carrying cell belongs to a clone significantly enriched for that variant, and non-informative otherwise (**Fig. 2C**). Much of the lineage signal came from calls with few supporting eUMIs and low per-cell heteroplasmy (mutant eUMIs / total eUMI depth). Among lineage-informative calls, 42.0% were supported by 1–3 eUMIs per cell and 52.9% fell below 10% per-cell heteroplasmy (**Fig. 2D**). These are therefore not negligible byproducts of technical noise but a substantial share of the recoverable signal. Against a reshuffled background, every eUMI and heteroplasmy group was significantly enriched for lineage-informative calls, and informativeness rose with molecular support — 87.7%, 89.0%, and 93.0% for calls supported by 2, 3, and ≥4 eUMIs (**Fig. 2E**). The 1-eUMI and lowest-heteroplasmy groups were the noisiest, yet still significantly exceeded the shuffled background and contributed to clonal recovery.

This benchmark rests on a lentiviral-barcode–anchored human HSC system. While this is a representative human primary system, optimal parameters and the relative performance of filtering strategies may vary across cell types, tissues and species. Also, the ground truth itself is imperfect: barcode silencing, recovery dropout, and mtDNA substructure may cause genuine lineage signals to be missed or misclassified, which likely makes our estimates of lineage informativeness conservative.

Finally, building a precise and confidence-aware lineage tree from mtDNA variants remains a related but distinct computational challenge and an active area of development [11–13]. Because mtDNA molecules are replicated and partitioned stochastically during cell division, heteroplasmy levels can drift across generations, creating phylogenetic uncertainty. Addressing this uncertainty requires inference frameworks that model mitochondrial inheritance, distinguish confident clonal relationships from ambiguous ones, and report interpretable measures of support. Drift-aware approaches such as MitoDrift provide one path toward this goal, extending mtDNA lineage tracing toward quantitatively calibrated lineage reconstruction [11].

## Conclusions

These analyses provide a ground-truth–anchored framework for best practice in mtDNA-based single-cell lineage tracing. We recommend redeemR2.0/filter2, a UMI-enabled error filtering approach, as the default preprocessing strategy: its broad, molecularly filtered variant spectrum gave the best clonal reconstruction of the strategies tested. Restrictive heteroplasmy thresholds carry a substantial sensitivity cost, since variants below 10% accounted for more than half of the lineage-informative signal and their inclusion improved both precision and recall. Calls with low molecular support, including single-eUMI calls, should likewise be retained when the goal is overall clonal structure. For variant-level interpretation, however, low-molecule calls should be evaluated cautiously: their value is collective rather than individual. Robust mtDNA lineage tracing thus benefits from pairing rigorous molecule-level filtering with inclusion across the full heteroplasmy spectrum.

## Methods

### Study design and materials

To benchmark lineage inference against an independent clonal reference, primary human HSPCs from a healthy donor were transduced with the LARRY v2 lentiviral barcode library, expanded ex vivo, and profiled by single-cell multiome sequencing with targeted capture of lentiviral barcodes and mtDNA (ReDeeM). Barcodes were assigned with BARtab, and clones of at least two cells defined the ground truth. The data was generated and shared publicly previously[11].

### Variant calling and filtering

REDEEM-V consensus calls were processed in redeemR2.0. filter2, its default post-consensus workflow, applies 9-bp fragment-edge trimming; retains cells with mean mtDNA coverage above 10 and variants seen in two or more cells; applies a per-variant binomial goodness-of-fit test against a technical-noise model (FDR 0.05); and removes homoplasmic, RSRS50 and blacklisted variants using MITOMAP population frequencies [14]. filter1 denotes the original redeemR strategy [3].

### Benchmarking

filter1, filter2, filter2 excluding 1-eUMI calls, filter2 at heteroplasmy thresholds of 0.02-0.10, and mgatk (fr2_mc3, at baseline and with a 0.07 heteroplasmy threshold [15,16]) were compared on ten fixed clone-based subsets of 200 cells, with one MitoDrift tree per condition-subset at matched parameters. Across MitoDrift support thresholds, internal nodes were treated as inferred clades, and recall and clade precision scored against lenti-clones at Jaccard 0.5 and 0.75, averaged across subsets.

### Statistical analysis

Variant-clone association was tested by two-sided Fisher’s exact test with Benjamini-Hochberg correction across clones (FDR 0.01). Background expectations came from a null model preserving each variant’s positive-cell number, eUMI support and depth while randomizing cell assignment.

### Software

The updated ReDeeM variant processing are included in R package REDEEM-R (redeemR2.0) (https://github.com/sankaranlab/redeemR). The reproducibility code are available through Figshare (https://figshare.com/s/682b71b1e62fda514692)

More detailed method descriptions are provided in supplementary methods.

## Data availability

Processed sequencing data of Lentiviral barcoding ground-truth datasets for benchmarking mtDNA-based lineage tracing has been shared previously (Gao et al. 2026) and is available on figshare (https://doi.org/10.6084/m9.figshare.31323481).

## Abbreviations

AUPRC: Area under the precision–recall curve
eUMI: Endogenous unique molecular identifier
FDR: False discovery rate
HSPC: Hematopoietic stem and progenitor cell
mtDNA: Mitochondrial DNA

## Ethics approval

Not applicable

## Acknowledgements

We thank members of the Sankaran and Weissman labs for valuable comments. J.S.W., and V.G.S. are investigators of the Howard Hughes Medical Institute.

## Authors’ contributions

C.W., J.S.W., and V.G.S. conceived the project and directed the studies. C.W. directed the experimental studies and computational analyses. T.G., J.G., I.J., M.P., assisted with data analysis. J.S.W. and V.G.S. supervised the study. All authors contributed to writing the manuscript.

## Supplementary Methods

### redeemR2.0 variant filtering and annotation

redeemR2.0 is the updated downstream analysis framework used to process REDEEM-V mitochondrial consensus variant calls before lineage reconstruction. The framework includes variant filtering, annotation, quality control, matrix construction, and utilities for downstream lineage analysis. In this manuscript, filter2 refers specifically to the default redeemR2.0 post-consensus variant-filtering workflow used to generate the filtered mtDNA variant set analyzed downstream. We use this term to distinguish the updated filtering strategy from the original redeemR filtering strategy, referred to as filter1(1). Filter2 is applied after UMI-based consensus variant calling and is designed to retain high-fidelity somatic mtDNA variants while removing residual technical artifacts, inherited or pre-existing polymorphisms, and variants less suitable for neutral lineage tracing.

REDEEM-V consensus variant calls were imported into redeemR. Unless otherwise specified, analyses used the stringent consensus output (S). To reduce residual fragment-end artifacts, raw genotype calls were annotated by their distance to the nearest fragment end, and calls falling within the trimmed edge window were removed. The default redeemR2.0 preprocessing used a 9-bp edge-trimming threshold. Genotype summaries were then reconstructed after trimming, including per-cell variant UMI counts, site depth, and heteroplasmy (calculated as mutant eUMIs divided by total site-covering eUMIs) for each retained cell–variant observation.

Initial redeemR objects were generated from the trimmed genotype summaries. Cells were retained if their mean mitochondrial coverage was at least 10. Candidate variants were initially required to be detected in at least two cells, with at least one supporting UMI in an individual cell. Homoplasmic or near-homoplasmic variants were annotated based on broad detection across covered cells, high mean heteroplasmy, and low variability, using the following criteria: CellNPCT > 0.75, PositiveMean > 0.75, and CV < 0.01 (CellNPCT denotes the fraction of covered cells in which the variant was detected, PositiveMean denotes the mean heteroplasmy among variant-positive cells, and CV denotes the variability of heteroplasmy across positive cells. Count matrices and binary cell-by-variant matrices were then constructed from heteroplasmic variants.

To distinguish recurrent biological signals from residual technical noise, redeemR2.0 applied a per-variant binomial goodness-of-fit test. For each variant, the distribution of variant UMI counts across cells was compared with a single-binomial technical-noise model parameterized by the observed total variant counts and position-specific mean coverage. A chi-square statistic was calculated from the observed and expected count frequencies, and p-values were converted to q-values. Variants were retained if they were heteroplasmic, detected in at least two cells, and passed an FDR threshold of 0.05.

Retained variants were then annotated for population, sequence-context, and functional features. These annotations included MITOMAP (https://www.mitomap.org/MITOMAP) population frequencies, RSRS50 ancestral-state status, haplogroup marker counts, mitochondrial blacklist regions (Mitochondrial blacklist regions were defined as recurrently problematic rCRS intervals prone to misalignment or homopolymer-associated artifacts, including positions 302–315, 513– 525, 3105–3109, and 16182–16187, homopolymer context, hypermutable-site labels, amino-acid consequence and predicted coding impact using a mitochondrial coding reference, mitochondrial disease annotations, and transition/transversion class. For downstream lineage analysis, redeemR2.0 removed heteroplasmic RSRS50 variants and variants falling within predefined mitochondrial blacklist regions. Because these annotation fields are retained in the output, users can apply additional study-specific filters based on population, sequence-context, or functional criteria as needed.

A depth matrix matched to the filtered cell-by-variant count matrix was then generated from QualifiedTotalCts (QualifiedTotalCts is the REDEEM-V per-cell, per-position mtDNA coverage table after consensus filtering). It was used to construct a depth matrix matched to the filtered cell-by-variant count matrix, enabling heteroplasmy calculation and depth-aware variant filtering. Median per-variant depth and the number of cells with nonzero coverage were added to the variant table. per-variant UMI-support features were added for downstream quality assessment, including the fraction of detections supported by two or three UMIs and the mean UMI count among detections supported by more than one UMI.

The final redeemR2.0 object contained filtered genotype summaries, variant annotations, cell-by-variant count and binary matrices, a matched depth matrix, and quality-control summaries. QC outputs included mutation-spectrum summaries, transition/transversion ratios, depth distributions, variant-support plots, matrix-dimension logs across filtering steps, and filter2 diagnostic plots.

### LARRY ground-truth benchmark

To benchmark mtDNA-based lineage reconstruction against a static ground truth, we used a lentiviral barcoding system (LARRY) in primary human HSCs coupled with joint scRNA-seq and scATAC-seq and deep mtDNA capture (ReDeeM)(1–3). Briefly, HSCs from a healthy donor were transduced with the LARRY lentiviral barcode library and plated for in vitro differentiation. Following ex vivo expansion, matched single cells were profiled with scRNA+scATAC multiome sequencing with targeted capture to read out (i) each cell’s LARRY barcode (defining its ground-truth clone membership) and (ii) mtDNA sequences used for MitoDrift lineage inference.

Ground-truth clonal identities defined by lentiviral barcodes were processed and annotated using BARtab (https://github.com/DaneVass/BARtab). We reconstructed redeemR lineage trees under two default preprocessing choices (“Filter1” and “Filter2”) and benchmarked them against curated ground-truth clones. For each ground-truth clone, we computed its maximum Jaccard similarity (Max Jaccard) to any inferred clade. Agreement was visualized with mutation frequency heatmap aligned to ReDeeM tree tip order side-by-side. The clone-wise Max Jaccard are shown for each clones by accuracy bars (Fig. 1E).

For the benchmarking, we compared lineage reconstruction precision-recall performance across predefined filtering strategies in ReDeeM and mgatk using a matched evaluation framework. For ReDeeM, we analyzed S consensus with Filter1 (keep consensus-called mutations that recur across cells and reach ≥2 eUMIs in at least one cell), redeemR2.0/filter2 (Filter1 + fragment edge-trimming + binomial/FDR test to remove remaining noise), Filter2 with no-1UMI (mutation observations with 1-UMI support are removed), and Filter2 with additional heteroplasmy thresholds (≥ 0.02, 0.04, 0.06, 0.08, 0.10); for mgatk, we used fr2_mc3 preprocessing (minimum 2 supporting reads per strand; variant must be detected in at least 3 confident cells; https://github.com/caleblareau/mgatk) with baseline or additional heteroplasmy thresholds ≥ 0.07 settings(4). Ground-truth cell-clone annotations were derived from curated barcode metadata (clone size ≥2). We generated 10 fixed clone-based subsets and reused the same subset definitions for every condition to control for composition effects, then inferred one MitoDrift tree per condition-subset with matched inference parameters (Fig. 2A-B).

We evaluated mtDNA-based lineage reconstruction using a MitoDrift confidence-sweep framework. Briefly, MitoDrift models single-cell mtDNA variant observations under mitochondrial heteroplasmy drift and measurement noise to infer lineage trees with branch-level support, allowing low-confidence branches to be progressively collapsed by varying a support threshold, τ. At each τ, we treated each internal node in the confidence-refined tree as an inferred clade and compared its cell membership to LARRY barcode-defined ground-truth clones using Jaccard overlap. Clone recall was defined as the fraction of ground-truth Lenti-clones recovered by at least one inferred clade above a specified Jaccard threshold, whereas clade precision was defined as the fraction of inferred clades matching a Lenti-clone above the same threshold. We evaluated performance across τ using fixed Jaccard thresholds of 0.5 and 0.75. In addition to end-to-end precision–recall, in which all conditions were evaluated on the same cell set, we also computed conditional precision by restricting the precision calculation to cells evaluatable after filtering, while retaining recall over the full cell set. Per-subset curves were averaged across ten matched 200-cell subsamples at each τ. Full details of the MitoDrift model and confidence-calibrated tree inference are described in the MitoDrift study(3).

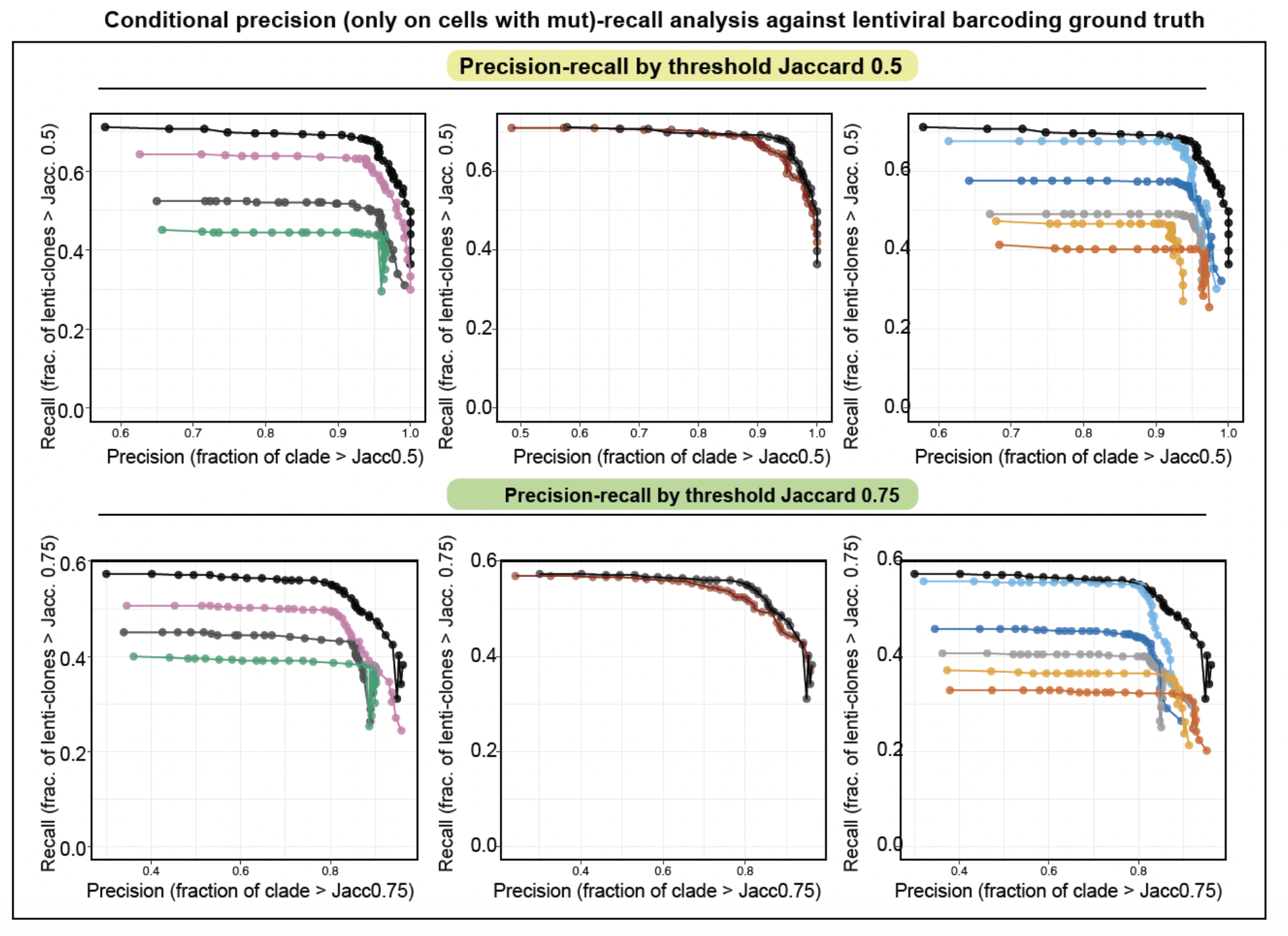

*Supplementary figure, conditional precision-recall analysis, related to Fig. 2B*

### Informative mtDNA variant-call analysis

To evaluate how lineage information was distributed across individual mtDNA mutation calls, we performed a per-variant, call-level analysis using LARRY barcode-defined clones as independent ground truth.

For each retained mtDNA variant, we identified all cells in which the variant was detected. A mutation call was defined as detection of a specific mtDNA variant in a specific cell. Variant-positive cells were linked to LARRY-defined clones using matched cell-barcode annotations. Variants detected in fewer than five LARRY-barcoded cells were excluded from this analysis.

We next tested whether each mtDNA variant was significantly associated with individual LARRY-defined clones. For each variant–clone pair, we constructed a 2 × 2 contingency table comparing variant-positive and variant-negative cells inside versus outside the clone and applied Fisher’s exact test. P values were adjusted across clones using false discovery rate correction. Clones with FDR-adjusted P ≤ 0.01 were considered significantly associated with that variant. A mutation call was labeled lineage-informative if the cell carrying the variant belonged to a clone significantly associated with that variant; calls outside such clones were labeled non-informative.

To quantify how informative calls were distributed across molecular-support levels, we summarized lineage-informative and non-informative mutation calls by the number of supporting eUMIs in each cell. Calls were first summarized in detailed eUMI bins of 1, 2, …, 9, and ≥10 eUMIs, and were additionally grouped into bins of 1, 2, 3, and ≥4 eUMIs for visualization. For each eUMI bin, the informative-call fraction was calculated as the number of lineage-informative calls divided by the total number of mutation calls in that bin. As a background control, we generated a random-shuffle null model. For each variant, the null preserved the number of variant-positive cells as well as the observed eUMI-support and depth values, but randomly reassigned those calls to barcode-annotated cells. The same variant inclusion criteria, call-level filtering, Fisher’s exact test, FDR threshold, and eUMI-bin summarization were then applied to the shuffled data. This provided the expected informative-call fraction under randomized cell–clone relationships and an empirical p value is derived.

